# Short duration overnight cattle kraaling in natural rangelands: does time after kraal use affect their utilization by wildlife and above ground grass parameters?

**DOI:** 10.1101/2021.03.08.434375

**Authors:** Rangarirai Huruba, Servious Nemera, Faith Ngute, Meshack Sahomba, Peter J. Mundy, Allan Sebata, Duncan N. MacFadyen

## Abstract

Short duration overnight cattle kraaling in natural rangelands creates nutrients hotspots attractive to a diverse suite of large mammalian herbivores. However, few studies have determined the use of these sites by large mammalian herbivores. We determined the number of animal sightings per day from camera traps and used them as proxies for use of these newly created nutrient hotspots of varying ages (1, 2, 3 and 4 years) and surrounding vegetation. Six mammalian herbivores of different sizes belonging to three feeding guilds *viz*. grazers (Burchell’s zebra *Equus quagga burchelli* and warthog *Phacochoerus africanus*), mixed feeders (impala *Aepyceros melampus* and African savanna elephant *Loxodonta africana africana*) and browsers (northern giraffe *Giraffa camelopardalis giraffa* and greater kudu *Tragelaphus strepsiceros*) frequently used these nutrient hotspots. The number of sightings per day of mammalian herbivores was determined during three periods of the year (January – wet season; June – early dry season; October – late dry season) to ascertain their use of these nutrient hotspots. In addition, above ground grass biomass and height was measured and related to grazer sightings. Furthermore, we tested if repeated grazing in the newly created nutrient hotspots stimulated grass compensatory growth. All the mammalian herbivores used the newly created nutrient hotspots similarly throughout the year, with impala the most active users. Grazer and browser use of nutrient hotspots was not influenced by their age, while mixed feeders mostly used the one year old sites. Grazer use of nutrient hotspots was not influenced by aboveground grass biomass and height. Repeated clipping (proxy for grazing) resulted in compensatory aboveground grass biomass growth in nutrient hotspots. Impala benefited the most and zebra the least from the creation of nutrient hotspots in natural rangelands. We conclude that creation of nutrients hotspots through short duration overnight kraaling results in rangeland heterogeneity that improves availability of herbivore foraging sites.

## Introduction

In African savanna ecosystems availability of nutritive forage for both domestic and wild herbivores is a major constraint to their production. Hence, both domestic and wild herbivores often congregate in termite mounds and grazing lawns which act as nutrient hotspots in search of nutritive forage [1]. The availability of termite mounds and grazing lawns results in spatial heterogeneity in rangelands, improving availability of nutritive forage for both domestic and wild ungulates, particularly in nutrient poor African savanna ecosystems. In addition to termite mounds and grazing lawns old cattle bomas or corrals (also referred to as glades) are increasingly being recognized as important nutrient hotspots in east and southern Africa [2-6]. When large herbivores congregate in old bomas for foraging and to seek refugee against predators [7], they alter the ecosystem functions through redistribution of nutrients within the terrestrial ecosystem [8]. Large herbivores harvest nutrients from the surrounding landscape through consumption of plant material and then deposit them in bomas as dung and urine overnight [9, 10]. The continuous use of old bomas by large mammalian herbivores keeps them nitrogen enriched and highly productive through dung and urine addition [11]. Nitrogen is mostly recycled as urine and phosphorus through dung deposition [12]. Some private ranch owners in east and southern Africa are now corralling (hereafter kraaling) cattle overnight in temporary mobile overnight kraals to simulate the nutrient hotspots created by old cattle bomas (see Huruba et al.[2, 3], Porensky and Veblen [5] and Sibanda et al. [4]). Few studies have monitored the use of nutrient hotspots created through kraaling cattle in temporary mobile overnight kraals by wildlife (but see Huruba et al. [2]).

Large mammalian herbivore behavioral ecology studies are now increasingly using camera traps [13, 14]. The use of nutrient hotspots by wildlife can be monitored using camera traps as they are cost-effective, efficient and non-intrusive [15]. Camera traps can be used to determine abundance of large mammalian herbivores and more specifically look at their spatial and temporal use of resources in ecosystems [16].

Nutrient hotspots are important foraging sites for wildlife [17], with species such as impala (*Aepyceros melampus*) regular users [18]. Wildlife is attracted to nutrient hotspots because of nutritious forage and openness that improves predator detection [7, 19]. Large herbivores such as Burchell’s zebra (*Equus quagga burchelli*), warthog (*Phacochoerus africanus*), African elephant (*Loxodonta africana africana*), giraffe (*Giraffa camelopardalis giraffa*) and greater kudu (*Tragelaphus strepsiceros*) have been observed in nutrient hotspots created through kraaling cattle in temporary mobile overnight kraals (Huruba pers. obs.). Classifying zebra, warthog, impala, elephants, giraffe and greater kudu into functional groups using their feeding guild could improve our understanding of their use of nutrient hotspots. In this study we classified the six herbivores into three feeding guilds *viz*. grazers (zebra and warthog), mixed feeders (impala and elephants), browsers (giraffe and greater kudu). Burchell’s zebra is a large grazer tolerant to fibrous diets, warthog is a small grazer intolerant of fibrous diets, impala is a highly selective medium size mixed feeder, African savanna elephant is a less selective large mixed feeder, while giraffe and greater kudu are large obligate browsers [20, 21].

The management of natural rangelands in a way that enables the creation of patchiness and heterogeneity is considered of benefit in improving plant biodiversity and productivity [22]. One such practice is setting up short duration overnight cattle kraals in natural rangelands [2, 3, 5]. Short duration overnight kraaling results in diversity in vegetation structure and function through creating patches of nutrient hotspots. The nutrient hotspots act as niches for wildlife to occupy improving the ability of rangelands to provide forage to herbivores even during times of shortages [23]. Repeated grazing in previously kraaled sites, in addition to urine and dung deposition helps to maintain grasses in a nutritive vegetative state [5]. The grazer-grass feedback loop ensures continued availability of nutritious forage for the herbivores [24]. Different suites of herbivores are expected to show variation in their use of previously kraaled sites. For example, small grazers maybe attracted to previously kraaled sites by nutritious grass regrowth [25], due to their small digestive systems which constrain their ability to digest fibrous diets [26], while large grazers are more tolerant to fibrous diets [27]. Therefore, large grazers, such as zebra, are expected to use previously kraaled sites less frequently than the surrounding vegetation because of their tolerance to poor quality forage [25, 27]. In addition, prey species may seek refuge from predators in previously kraaled sites which are more open than the surrounding vegetation [7].

Utilization of nutrient hotspots by large herbivores is expected to vary with time of year, particularly in African savanna ecosystems which are characterized by distinct wet and dry seasons with different plant growth. Grasses are green and nutritious during wet season but brown and less nutritious during dry season. The change in nutritive value of grass between wet and dry season is expected to influence the use of nutrient hotspots. However, repeated grazing of nutrient hotspots could stimulate grass regrowth even during the dry season if there is adequate soil moisture leading to the production of nutritive forage. During wet season green grass is widespread in the rangeland affording herbivores a wider choice of nutritive forage. In southern Africa the wet season occurs between November and April, while the dry season is between May and October. Mayengo et al. [1] reported grazing lawns as mostly used during dry season presumably because they are the only patches with green grass. Thus, the use of nutrient hotspots created through short duration overnight cattle kraaling by wildlife is expected to vary with the time of the year.

Soil nutrients that accumulate in nutrient hotspots created through short duration overnight cattle kraaling declines with time after kraal use [2, 3, 4]. The loss of soil nutrients with time after kraal use is expected to result in a decrease in grass quality leading to a decline in their utilisation by grazers. Grass nutrient influences the selection of foraging patches by grazers [28]. Thus, grazer use of previously kraaled sites is expected to be high one to two years after kraal use and then subsequently decline with time. However, repeated use of previously kraaled sites by grazers results in deposition of dung and urine enhancing or maintaining soil fertility. Dung and urine deposition in kraaled sites results in recycling of plant nutrients [29]. The influence of time after kraal use on the utilisation of these newly created nutrient hotspots by grazers has not been studied.

Relative abundance indices derived from camera traps show strong positive correlation to wildlife population estimates [30]. Thus, the probability of sighting animals in camera traps is strongly influenced by their population. Therefore, the utilization of nutrient hotspots by wildlife could be influenced by animal abundance. To ascertain if use of nutrient hotspots is not influenced by animal abundance there is need to relate sighting index to population.

Nutrient hotspots created through short duration overnight kraaling are rich in soil nutrients resulting in accumulation of large aboveground grass biomass and growth of tall grasses. However, since these nutrient hotspots are attractive to grazers, persistent grazing limits aboveground grass biomass accumulation and restricts grass height. Muchiru et al. [31] reported densities of grazers as positively correlated to herbaceous biomass. Young et al. [32] reported grazers as foraging in grazing patches with high grass biomass to meet their nutrient requirements. Interestingly, the low aboveground grass biomass due to repeated grazing in nutrient hotspots could benefit herbivores such as impala. Impalas select short, low fiber grass which is highly digestible to increase forage intake [32]. Therefore, impalas are expected to benefit most from creation of nutrient hotspots because of short duration overnight kraaling. On the contrary, zebras are expected to be the least beneficiaries of short duration overnight kraaling since they select foraging patches with high aboveground grass biomass to achieve high digestive fill, because as hindgut fermenters they have a fast digesta passage rate [33]. However, zebra and other equids consume both short and tall grass to balance between forage quality and quantity [34]. Zebra have narrow muzzle considered well suited for clipping tall grasses [35]. Grass height influences herbivore habitat use [36].

Repeated grazing is thought to create a positive herbivore-grass feedback loop that maintains high soil and plant nutrient levels [37]. Grass regrowth in response to grazing is rich in nutrients and attracts grazers to nutrient hotspots [3, 38]. However, the positive herbivore-grass feedback has not been widely tested, particularly in nutrient hotspots. The positive effects of repeated grazing can easily be upset because of overgrazing that result in rangeland degradation [39]. Although repeated grazing or defoliation generally leads to increased relative grass growth rate [40], grasses could respond through under-, partial- or over-compensation of lost foliar tissue [38]. However, most grasses under or equally- compensate the lost biomass because defoliation results in loss of photosynthetic material limiting the ability of the plants to photosynthesize [41]. In addition, grass regrowth in nutrient hotspots could benefit from enhanced soil fertility due to dung and urine deposition [42]. In this study we investigated the effect of repeated grazing on aboveground grass biomass. Enhanced soil fertility in nutrient hotspots further improves the quality of foliage which is maintained through nutrient cycling via urine and dung deposition [43].

We studied the use of previously kraaled sites of varying ages by six mammalian herbivores of different sizes and feeding guilds during three periods of the year. We collated the number of large herbivore sightings in previously kraaled sites from camera trap photographs during three periods of the year (January – wet season; June – early dry season; October – late dry season) to ascertain their use of these nutrient hotspots. In addition, above ground grass biomass and height were measured and related to grazer sightings. Furthermore, we carried out an experiment on simulated grazing to determine the response of grass growth to repeated (three times during growth season) defoliation. We tested the following research hypotheses: 1) time of year does not affect use of previously kraaled sites by wildlife, 2) age of previously kraaled site does not affect its use by wildlife, 3) the sighting of wildlife is not influenced by its population, 4) aboveground grass biomass and height does not influence grazer use of previously kraaled sites, and 5) repeated grazing result in compensatory aboveground grass biomass growth in previously kraaled sites.

## Materials & method

### Study site

Debshan ranch is located in central Zimbabwe (29°13′E, 19°36′S; 1230m elevation) (Figure 1). It is a mixed cattle-wildlife ranch that covers an area of 800 km^2^ and supports a diversity of large mammal species that include impala (*Aepyceros melampus*), Burchell’s zebra (*Equus quagga burchelli*), warthog (*Phacochoerus africanus*), African savanna elephant (*Loxodonta africana africana*), northern giraffe (*Giraffa camelopardalis giraffa*) and greater kudu (*Tragelaphus strepsiceros*). The study area is characterized by a catenal vegetation pattern, with most areas consisting of grassed bushland with patches of Miombo woodland [44]. The dominant woody species is *Acacia karroo* Hayne with the major grass species being *Hyparrhenia filipendula* (Hochst.) Stapf., *Eragrostis curvula* (Schrad.) Nees., *Heteropogon contortus* (L.) Roem. & Schult., *Bothriochloa insculpta* (Hochst. Ex A. Rich.), *Digitaria milanjiana* (Rendle) Stapf., and *Panicum maximum* Jacq. Mean annual rainfall is 612 mm, with a rainy season that runs from November to March and a dry season from April to October [44]. Mean annual temperature is 22.6 °C, with October (31.4 °C) the hottest month and July the coldest (8.5 °C).

**Figure 1.**
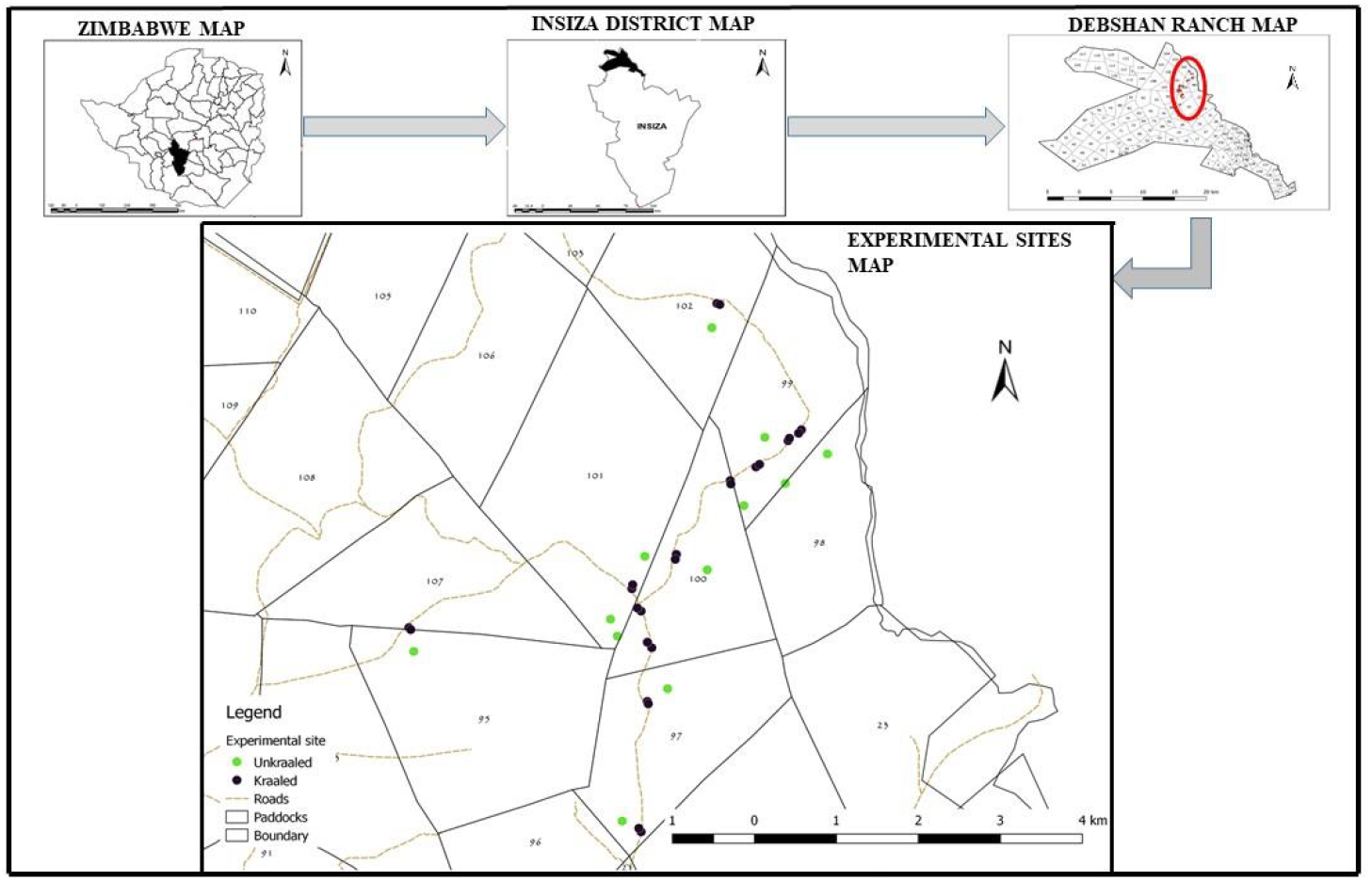
Debshan ranch location showing camera trap positions

### Camera traps setting

We deployed Cuddeback Attack/Attack IR digital scouting cameras (*n* = 11), Cuddeback C (modular) and E model cameras (*n* = 25) (Cuddeback Trail Camera company, India) infrared camera traps at eighteen locations in previously kraaled sites of varying ages (1, 2, 3 and 4 years) and control sites in surrounding vegetation at Debshan ranch between January and October 2017 (Fig. 1). The cameras were mounted on tree trunks at one meter above the ground to detect medium- to large-bodied mammals [45]. The cameras were programmed for pictorial (single capture/minute) data capture for diurnal and nocturnal animals at a trigger speed interval of 60 seconds as passive infrared sensors are triggered by motion or heat during the day or night. Date (dd/mm/yy), time (hh:mm) and camera number (ID) were recorded. Secure Digital (SD) memory cards and non-rechargeable batteries were replaced at two weeks interval. The pictorial data was downloaded from the SD cards and stored in folders labeled according to kraal age. Microsoft Excel version 2016 was used to store the photographic data with the following details: camera location (kraaled or unkraaled area), camera unit identifier, date (dd/mm/yy), time (hours, minutes) and animal species. Data collection was done in January (*n* = 30 days), June (*n* = 30 days) and October (*n* = 30 days) 2017. Camera trapping is non-intrusive and effective in studying mammal spatial and temporal use of habitats [46]. The number of animal sightings of each mammalian species during each period was recorded from the camera trap data and expressed as number of animal sightings per day.

### Estimates of aboveground grass biomass cropped by grazers

We set up four chicken wire mesh (2 cm diameter holes) herbivore exclusion movable cages (1m × 1m × 1 m) in each previously kraaled site and surrounding vegetation to estimate aboveground grass biomass cropped by grazing herbivores. The cages were kept in the same position during the growth season (October 2016 to May 2017). The difference in aboveground grass biomass inside and outside the movable cages was assumed to represent grass cropped by the grazing herbivores [47]. Aboveground grass biomass both inside and outside the movable cages was clipped using a clipper, air dried, before oven drying at 60°C for 48 hours and then weighed. Cropped aboveground grass biomass was then calculated as the difference between aboveground grass biomass inside and outside the mobile cages. Grass height was also measured in each sampling site using a tape measure.

To test if repeated grazing results in compensatory aboveground grass biomass growth we clipped grass inside movable cages three times during the growth season and compared with aboveground grass biomass in a single clipping. Aboveground grass biomass inside movable cages located in previously kraaled sites of different ages were clipped three times at twenty-eight days intervals from the beginning of the growing season (28/01/2017) (1st clipping), peak of the growing season (25/02/2017) (2nd clipping) and at the end of the growing season (25/03/2017) (3rd clipping). The aboveground grass biomass removed at each of the three clippings was recorded and the sum for all the clipping calculated. Compensatory aboveground grass biomass (gm^-2^) was calculated using the formula: total aboveground grass biomass for all three clippings – aboveground grass biomass clipped once at the end of growth season.

### Statistical analysis

A total of 2833 camera images captured during the study period (90 days) were used for analysis of number of wildlife sightings (January: *n* = 324; June: *n* = 874; October: *n* = 1635).

All data was tested for homogeneity of variance using the Levene statistics in IBM SPSS 16 prior to statistical analyses and normality using the Shapiro-Wilk test. First a general linear model univariate was used to test for the effects of wildlife species and age of kraal (time after kraal use) on animal sightings per day. Then the effects of time of year and age of kraal on use of previously kraaled sites were tested using one way analysis of variance and Tukey’s HSD used to determine *post hoc* differences within those effects.

## Results

Zebra, warthog, impala, elephants, giraffe and greater kudu were the most frequently sighted in the camera traps.

The results from the general linear model univariate analysis showed that overall, time after kraal use had no significant effect on wildlife sightings (*F*_1,4_ = 1.24, *p* = 0.32), while wildlife sighting varied with wildlife species (*F*_5,20_ = 7.30, *p* < 0.001). Time after kraal use x wildlife species interaction was significant (*F*_20,55_ = 9.00, *p* < 0.001).

The results from one way analysis of variance showed that wildlife sightings were similar in January, June and October for all species (zebra: *F*_2,9_ = 1.45, *p* = 0.285; warthog: *F*_2,9_ = 2.40, *p* = 0.146; impala: *F*_2,9_ = 0.72, *p* = 0.511; elephant: *F*_2,9_ = 0.02, *p* = 0.899; giraffe: *F*_2,9_ = 0.08, *p* = 0.926; kudu: *F*_2,9_ = 0.24, *p* = 0.795), with impala sightings higher than the other five wildlife species during the three times of the year (January: *F*_4,15_ = 52.07, *p* < 0.001; June: *F*_5,18_ = 3.87, *p* = 0.02; October: *F*_5,18_ = 4.72, *p* = 0.01) (Figure 2).

**Figure 2.**
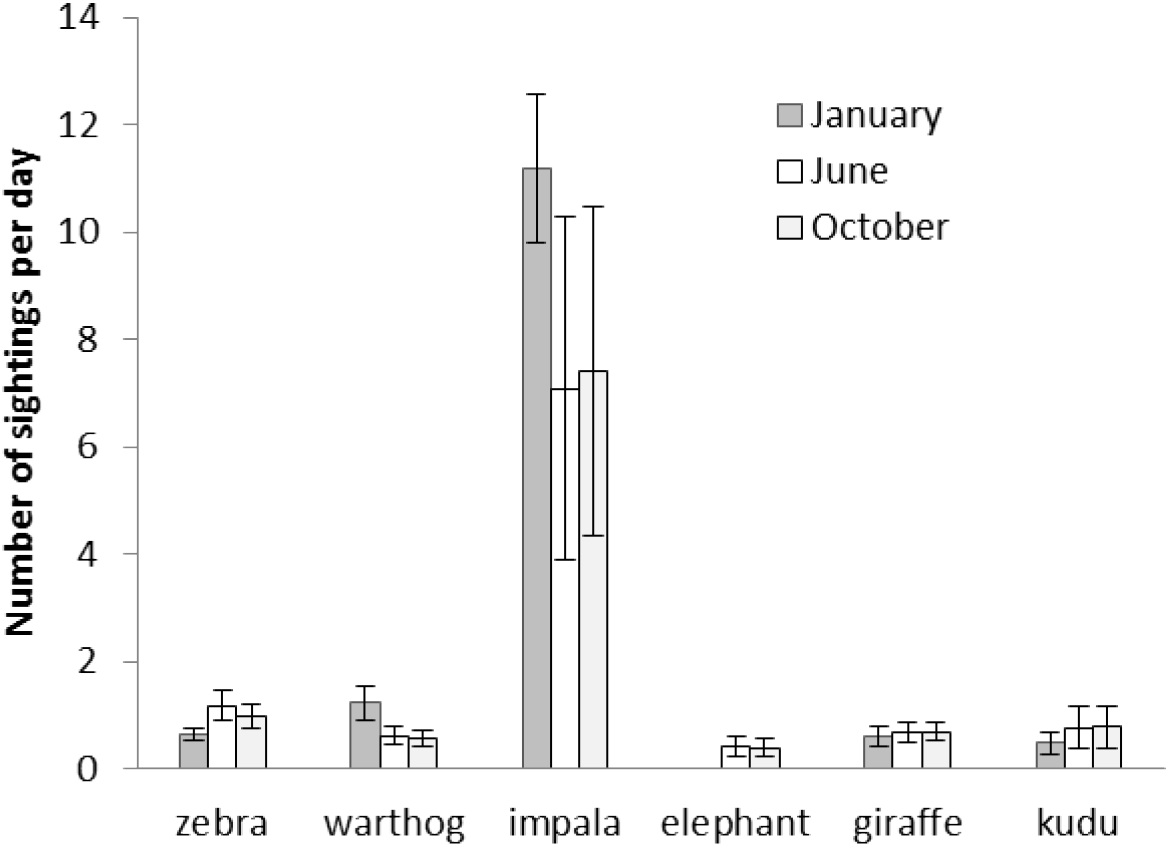
Mean (±SE) number of wildlife sightings per day during three periods (January, June and October) in previously kraaled sites

One way analysis of variance results showed that the number of wildlife sightings varied with age of kraal (time after use) in impala (*F*_4,10_ = 11.06, *p* = 0.001) and elephants (*F*_4,10_ = 153.45, *p* < 0.001) but not in the other wildlife species (Burchell’s zebra: *F*_4,10_ = 2.80, *p* = 0.09; warthog: *F*_4,10_ = 1.66, *p* = 0.24; giraffe: *F*_4,10_ = 2.11, *p* = 0.15 and greater kudu: *F*_4,10_ = 2.39, *p* = 0.12) (Figure 3). Impala mostly used previously kraaled sites one and four years after use, while African savanna elephant (*Loxodonta africana africana*), mostly preferred to use previously kraaled sites one year after use. Number of wildlife sightings within kraaling treatment varied with species (no kraaling: *F*_5,11_ = 3.95, *p* = 0.03; one year after kraaling: *F*_5,11_= 43.88, *p* < 0.001; three years after kraaling: *F*_5,11_ = 7.68, *p* = 0.002; four years after kraaling: *F*_5,11_ = 143.67, *p* < 0.001), except in the two years after kraaling treatment (*F*_5,11_ = 0.90, *p* = 0.517). Impala were the most active users of previously kraaled sites.

**Figure 3.**
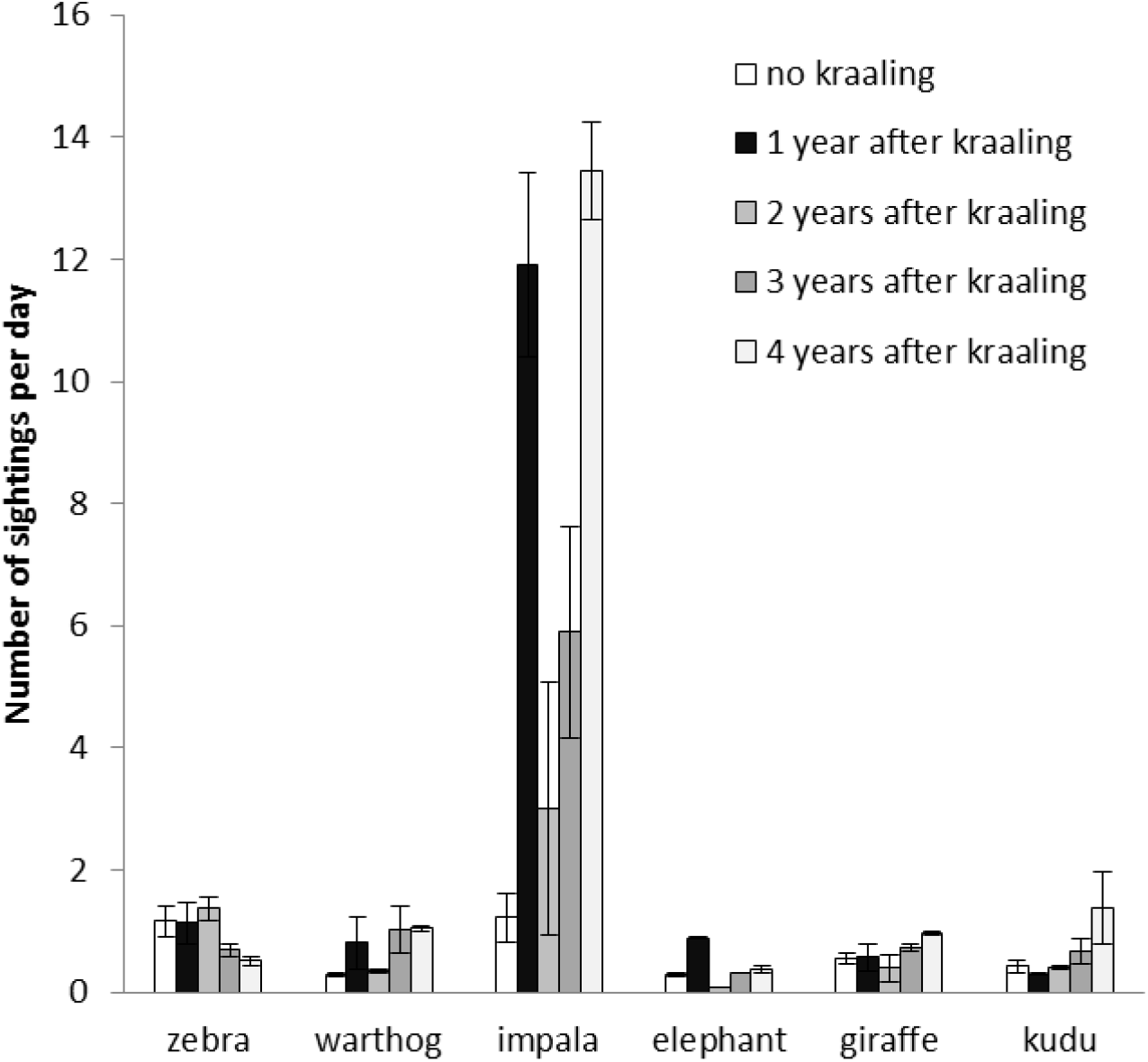
Mean (±SE) number of wildlife sightings per day in previously kraaled sites of varying ages

Impala and giraffe were the most and least abundant wildlife species at Debshan ranch in 2016 respectively (Figure 4). Giraffe had the highest sighting index (*F*_5,18_ = 7.02, *p* = 0.001; Figure 5). Sighting index was not significantly correlated to wildlife population (*r* = 0.10, *p* = 0.85, *n* = 6).

**Figure 4.**
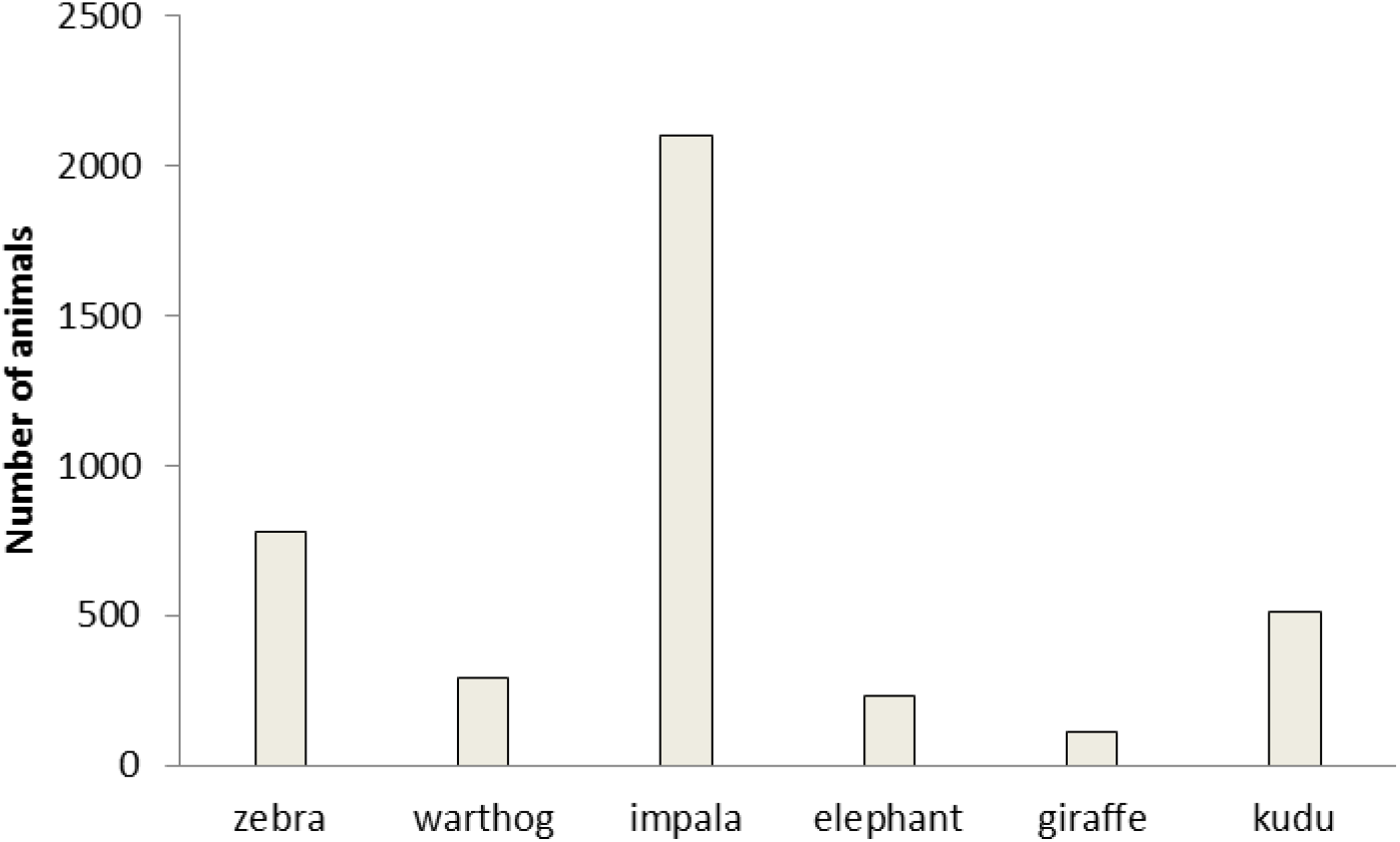
Population of six wildlife species at Debshan ranch in 2016

**Figure 5.**
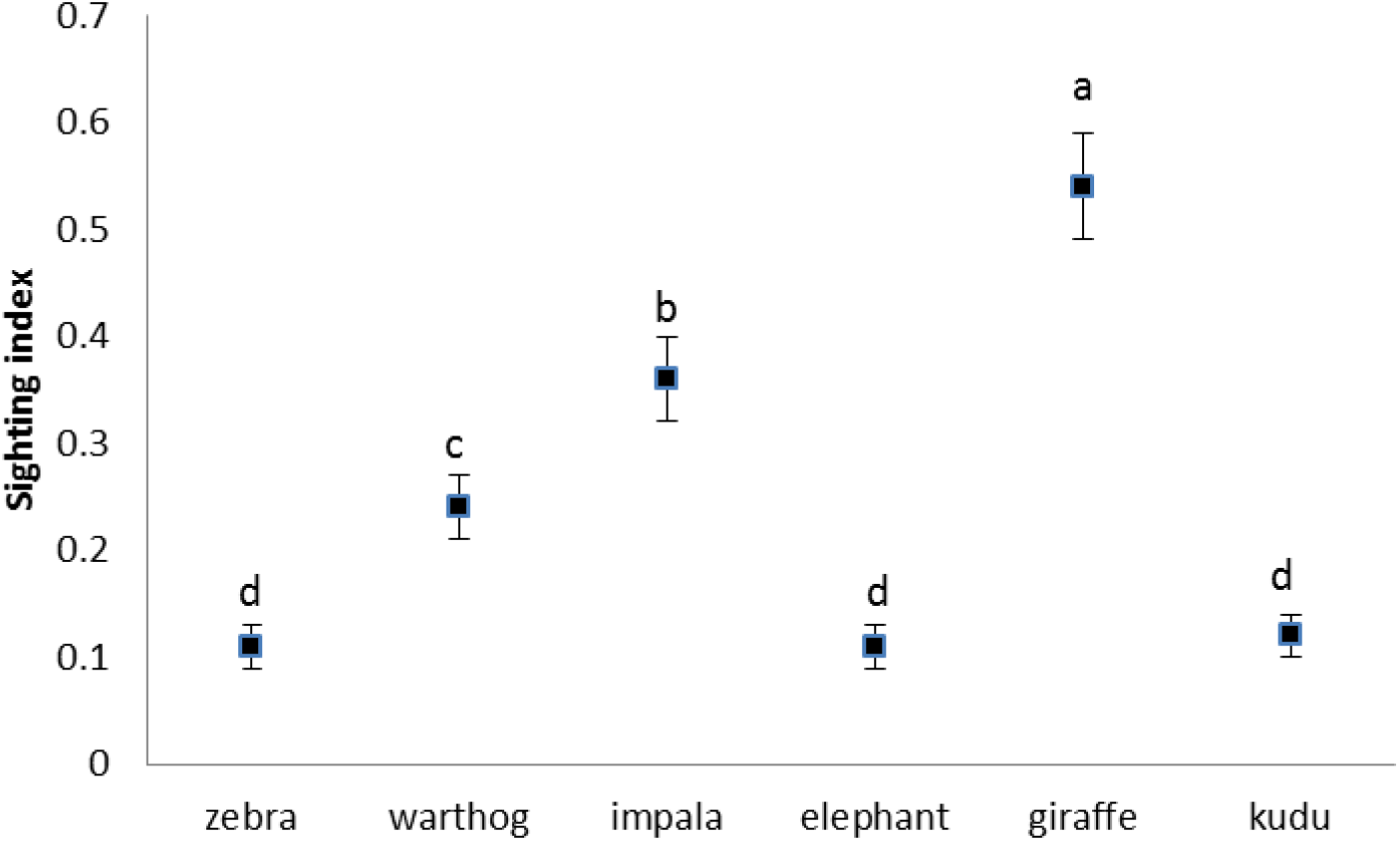
Mean (±SE) sighting index for six wildlife species in previously kraaled sites

Aboveground grass biomass was highest in surrounding vegetation (unkraaled sites) (*F*_4,20_ = 1167, *p* < 0.001), whilst among previously kraaled sites it was highest in the one year old sites and thereafter declined with age (Figure 6). The highest aboveground grass biomass was cropped in three year old previously kraaled sites (*F*_4,20_ = 112.98, *p* < 0.001; Figure 6). Grazing mostly occurred in the three and four year old previously kraaled sites with little taking place in the surrounding vegetation and the one year old previously kraaled sites. Grass was tallest in surrounding vegetation (unkraaled sites) (*F*_4,20_ = 407.13, *p* < 0.001; Figure 7), whilst among the kraaled sites it was tallest one year after kraal use and thereafter declined with time after use. Zebra and warthog sightings were not correlated to aboveground grass biomass (zebra: *r* = 0.54, *p* = 0.34, *n* = 5; warthog: *r* = - 0.68, *p* = 0.21, *n* = 5) and grass height (zebra: *r* = 0.57, *p* = 0.31, *n* = 5; warthog: *r* = - 0.76, *p* = 0.14, *n* = 5).

**Figure 6.**
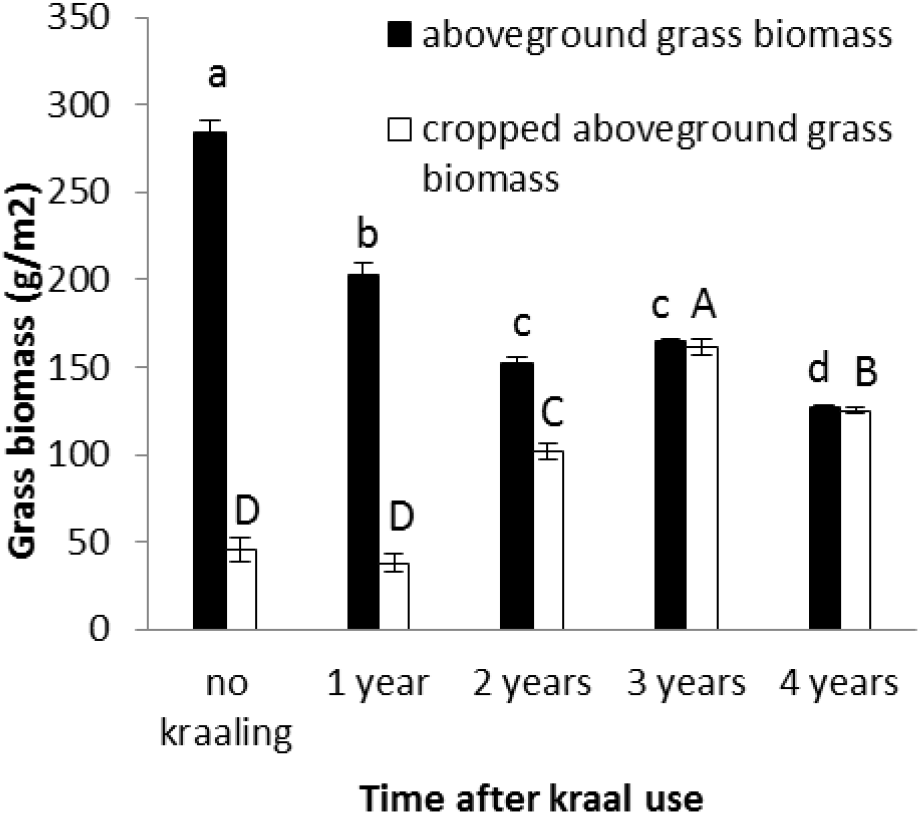
Mean (±SE) aboveground and cropped grass biomass at previously kraaled sites of different ages. Different letters (a, b, c and d - for aboveground grass biomass; and A, B, C, D - for cropped aboveground grass biomass) show differences in the treatments.

**Figure 7.**
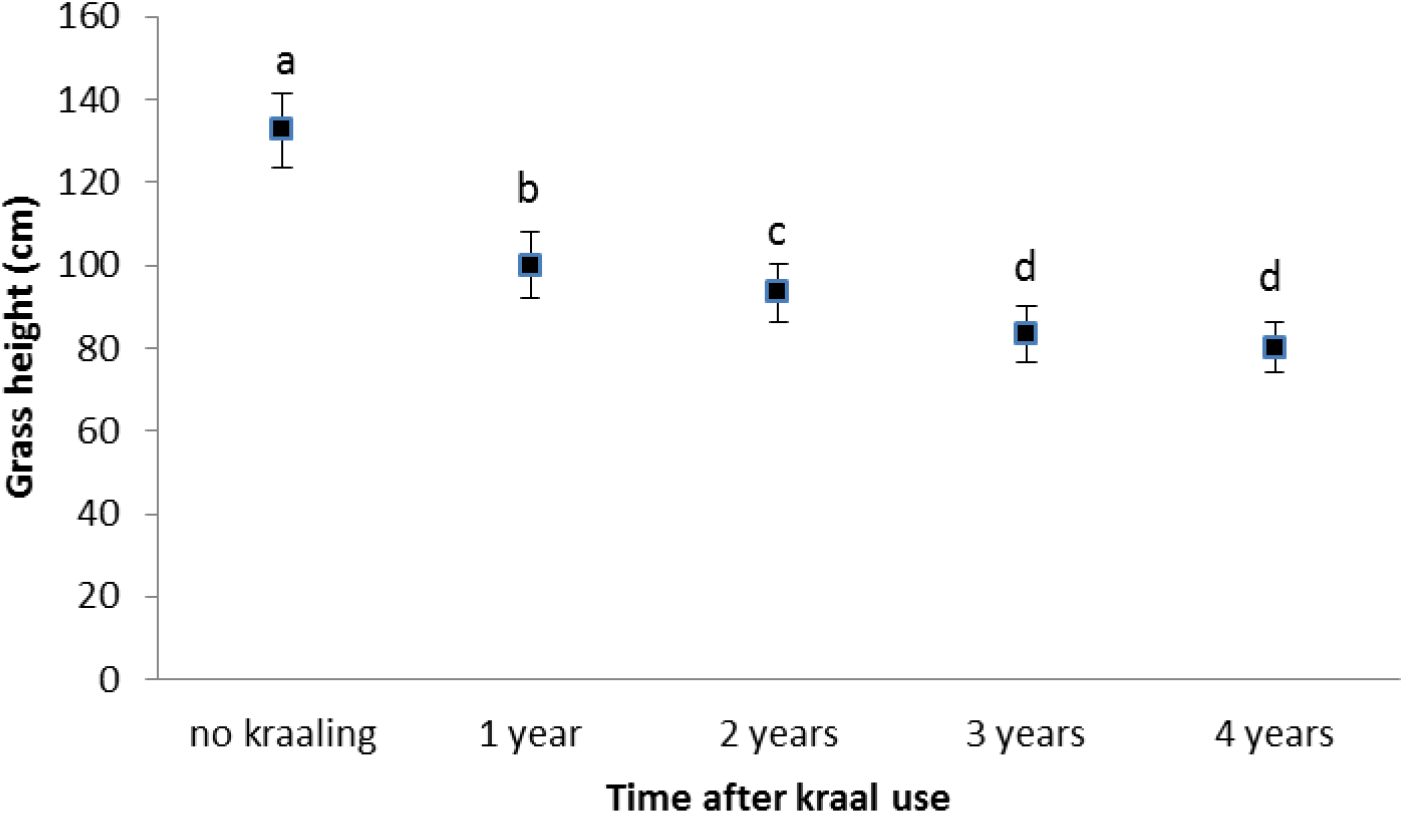
Mean (±SE) grass height in previously different ages

Repeated grass clipping (proxy for grazing) resulted in compensatory grass growth, with the highest in the one year after kraal use sites (Table 1).

**Table 1.** Mean (±SE) aboveground grass biomass (gm^-2^) in cages clipped once and repeatedly (three times) in previously kraaled sites of different ages

|  | Clipped once | Repeated clipping |  |  |  | Compensatory growth |
| --- | --- | --- | --- | --- | --- | --- |
|  |  | 1<sup>st</sup> clipping | 2<sup>nd</sup> clipping | 3<sup>rd</sup> clipping | Total for repeated clipping |  |
| 1 year after kraal use | 248.40<sup>b</sup> ± 4.46 | 379.20<sup>a</sup> ± 2.85 | 133.80<sup>b</sup> ± 1.69 | 85.20<sup>a</sup> ± 1.77 | 599.20<sup>a</sup> ± 2.08 | 351.20<sup>a</sup> ± 3.31 |
| 2 years after kraal use | 252.40<sup>b</sup> ± 3.75 | 331.20<sup>b</sup> ± 15.99 | 155.40<sup>a</sup> ± 1.29 | 72.60<sup>b</sup> ± 0.93 | 535.20<sup>b</sup> ± 1.83 | 274.20<sup>b</sup> ± 1.66 |
| 3 years after kraal use | 287.40<sup>a</sup> ± 3.33 | 281.60<sup>c</sup> ± 2.20 | 131.60<sup>b</sup> ± 1.62 | 71.05<sup>b</sup> ± 2.52 | 497.40<sup>c</sup> ± 1.81 | 209.40<sup>c</sup> ± 2.58 |
| 4 years after kraal use | 255.20<sup>b</sup> ± 4.19 | 205.00<sup>d</sup> ± 1.61 | 112.00<sup>c</sup> ± 0.95 | 37.00<sup>c</sup> ± 2.10 | 353.00<sup>d</sup> ± 4.36 | 101.00<sup>d</sup> ± 1.58 |
| | $F_{3,16} = 20.54$,<br>$p < 0.001$ | $F_{3,16} = 81.55$,<br>$p < 0.001$ | $F_{3,16} = 152.58$,<br>$p < 0.001$ | $F_{3,16} = 116.88$,<br>$p < 0.001$ | $F_{3,16} = 1453.00$,<br>$p < 0.001$ | $F_{3,16} = 1964.00$,<br>$p < 0.001$ |

## Discussion

Our research highlights the importance of short duration overnight cattle kraaling in creating nutrient hotspots for wildlife use in an African savanna ecosystem. We found that six mammalian herbivores belonging to three feeding guilds *viz*. grazers, mixed feeders and browsers frequently used nutrient hotspots created through short duration overnight cattle kraaling. We used the number of animal sightings per day from camera traps as a proxy for use of previously kraaled sites and surrounding vegetation. Bailey et al. [48] suggested that herbivores were able to identify patches with forage of varying nutritive value within the rangeland. Impala benefited the most and zebra the least from the creation of nutrient hotspots in the natural rangelands. Previous studies reported impala, warthog, elephants and other large herbivores also using nutrient hotspots created through overnight cattle kraaling [2, 4, 6].

Our findings supported the first hypothesis with wildlife use of previously kraaled sites similar throughout the year. This suggests that the newly created nutrient hotspots provide nutritive forage and / or refugee to wildlife throughout the year. As stated above impala were the most active users of these newly created nutrient hotspots throughout the year. Large herbivores actively seek nutrient hotspots in search of nutritive forage [2, 49]. Veblen and Porensky [6] reported giraffe as avoiding previously kraaled sites that had no woody plants, instead foraging in the surrounding vegetation with abundant woody species. Interestingly, in our study giraffe used previously kraaled sites more than the surrounding vegetation presumably because they were wooded. Our overnight cattle kraaling system did not cut down trees and shrubs, with plant damage due to trampling. The previously kraaled sites had both grass and browse, with trampling stimulating woody plant resprouting to produce nutritive browse. Traditional glades, formed from long duration kraaling, for example one year, in east Africa are treeless because trees are cut down for use as kraal fences [24]. In agreement with our findings, Veblen and Porensky [6] reported zebra as not actively seeking out high quality forage in nutrient hotspots presumably because their large size and hind-gut fermentation allowed them to consume fibrous diets. In addition, low aboveground grass biomass in previously kraaled sites made them less attractive to zebras [5].

Our results agreed with the second hypothesis in the case of grazers and browsers but not for mixed feeders. Grazer and browser use of previously kraaled sites was not influenced by their age. However, use by mixed feeders (impala and elephants) varied with age of previously kraaled sites. Both impala and elephants mostly used previously kraaled sites one year after use. In addition, impala also used previously kraaled sites four years after use frequently. Impala and elephants can switch between grasses and browse depending on their nutritive quality. Therefore, impala and elephants were able to make choices from previously kraaled sites of varying ages depending on the quality of either grasses or browse. An alternative explanation is that one year after use previously kraaled sites are generally open offering clear visibility to impala and unhinged movement of elephants. Other researchers also reported nutrient hotspots created from overnight kraaling as most attractive to mixed feeders such as impala and elephants [29, 50, 51]. Shannon et al. [51] attributed this to the ability of mixed feeders (particularly elephants) to mainly browse while also consuming grasses. Huruba et al. [2] reported cattle as breaking woody plant stems and stripping them of foliage initiating resprouting during overnight kraaling. The resprouts are attractive to herbivores due to high foliar nitrogen and low condensed tannin concentrations.

Our findings supported the third hypothesis that wildlife sighting was not influenced by its population. The high sighting index despite its low population suggests that giraffe were frequent users of the previously kraaled sites. Interestingly, impala had both high sighting index and population. This means that impala frequently used nutrient hotspots and had high populations.

Our results supported the fourth hypothesis with zebra and warthog use of previously kraaled sites not influenced by aboveground grass biomass and height. Although aboveground grass biomass and height were highest in surrounding vegetation this did not act as an attractant to the grazers, presumably, because the grass was moribund and thus unattractive to the grazers. In addition, areas with high aboveground grass biomass have high risks of predation from ambush-hunting predators such as lions [52]. Therefore, low aboveground grass biomass likely represents both preferred forage and low risk habitat for grazers. Further evidence for poor utilization of surrounding vegetation is shown by lower cropping as compared to three and four year old previously kraaled sites. Repeated grazing, particularly, by warthogs kept aboveground biomass and grass height low in previously kraaled sites especially the three and four year old previously kraaled sites. Huruba et al. [3] reported warthogs as intensely grazing in previously kraaled sites. Aboveground grass biomass cropped in all kraaled sites and surrounding vegetation were within the range of 89 to 951 gm^-2^ reported in the Kruger national park by Burkepile et al. [53]. Zebra can consume low quality grass because of their fast passage rate of forage through the gastrointestinal tract [26]. Zebras prefer grass with high biomass [54]. Groom and Harris [53] argued that forage quantity was more important than quality in sustaining large herbivores in resource-poor environments. Zebra do not show preference for grass of any height [55].

Our findings supported the fifth hypothesis with repeated clipping (proxy for grazing) resulting in compensatory aboveground grass biomass growth in previously kraaled sites. McNaughton [37] reported grasses in the Serengeti as over-compensating lost foliage. In addition, nutrient addition in the form of dung and urine could have enhanced grass growth in response to the grazing stimulus. Venter et al. [56] reported nutrient addition (in the form of animal dung) as increasing aboveground grass biomass. The decline in aboveground grass biomass regrowth between first and third clipping (see Table 3) was presumably due to resource exhaustion as a result of multiple grass resprouting in response to defoliation [57]. Mudongo et al. [58] also reported a decrease in aboveground grass biomass regrowth with increasing clipping frequency. In the long-term repeated grazing could negatively affect tillering leading to the loss of the grasses [59]. While previous studies have shown that grazing in the previous growing season reduces grass productivity in the next growing season [60, 61], this study shows that a decline in regrowth occurs in the current season. In addition, the decline in aboveground grass biomass regrowth with repeated clipping could be attributed to reduced soil moisture availability with advancing growth season which negatively affects nutrient mineralization [62]. Soil mineralization is higher early in the growing season when soil moisture is high as compared to late in the dry season [58].

## Conclusion

Our findings showed that short duration overnight cattle kraaling in natural rangelands creates nutrient hotspots which are attractive to wildlife. Grazers maintain a low above ground grass biomass and height through repeated grazing. Grass responds to repeated grazing through compensatory growth. More broadly, our results show that creation of nutrient hotspots through short duration overnight kraaling results in rangeland heterogeneity that improves availability of herbivore foraging sites.

## Acknowledgements

We wish to thank Debshan ranch owners and management for permission to carry out the study and for the financial support. We also wish to thank the associate editor and two anonymous reviewers for constructive and insightful contributions to this manuscript.

## Author Contributions

**Conceptualization:** Rangarirai Huruba, Allan Sebata

**Data curation:** Rangarirai Huruba

**Funding acquisition:** Rangarirai Huruba, Duncan N. MacFadyen

**Investigation:** Rangarirai Huruba, Servious Nemera, Faith Ngute, Meshack Sahomba

**Methodology:** Rangarirai Huruba, Allan Sebata

**Project administration:** Rangarirai Huruba

**Supervision:** Peter J. Mundy, Allan Sebata, Duncan N. MacFadyen

**Writing – original draft:** Rangarirai Huruba

**Writing – review & editing:** Rangarirai Huruba, Servious Nemera, Faith Ngute, Meshack Sahomba, Peter J. Mundy, Allan Sebata, Duncan N. MacFadyen

